# Default mode network anatomy and function is linked to pediatric concussion recovery

**DOI:** 10.1101/795740

**Authors:** Kartik K. Iyer, Andrew Zalesky, Karen M. Barlow, Luca Cocchi

## Abstract

**Objective:** To determine whether anatomical and functional brain features relate to key persistent post-concussion symptoms (PPCS) in children recovering from mild traumatic brain injuries (mTBI), and whether such brain indices can predict individual recovery from PPCS.

**Methods:** 110 children with mixed recovery following mTBI were seen at the concussion clinic at Neurology department Alberta Children’s Hospital. The primary outcome was the Post-Concussion Symptom Inventory (PCSI, parent proxy). Sleep disturbance scores (PCSI sub-domain) and the Neurocognition Index (CNS Vital Signs) were also measured longitudinally. PPCS was assessed at 4 weeks post-injury and 8-10 weeks post-injury. Grey matter volumes were assessed using magnetic resonance imaging (MRI) and voxel-based morphometry. Functional connectivity was estimated using resting-state MRI. Two complementary machine learning methods were used to assess if the combination of grey matter and functional connectivity indices carried meaningful prognostic information.

**Results:** Higher scores on a composite index of sleep disturbance, including fatigue, were associated with converging decreases in grey matter volume and local functional connectivity in two key nodes of the default mode network: the posterior cingulate cortex and the medial prefrontal cortex. Sleep-related disturbances also significantly correlated with reductions in functional connectivity between these brain regions. The combination of structural and functional brain indices associated to individual variations in the default mode network accurately predicted clinical outcomes at follow-up (area under the curve = 0.86).

**Interpretation:** These results highlight that the function-structure profile of core default mode regions underpins sleep-related problems following mTBI and carries meaningful prognostic information for pediatric concussion recovery.

## INTRODUCTION

Incidence rates of traumatic brain injury (TBI) in pediatric populations are on the increase, with median estimates suggesting 691 injuries per 100,000 emergency admissions ^1^. Most of these injuries are mild TBIs (∼90%, mTBI). A large proportion of children have persistent post-concussion symptoms (PPCS) following mTBI ^2, 3^ which is defined as the presence of at least two or more post-concussive symptoms that persist for 4 weeks or longer ^4^. Around 12% of children sustaining a mTBI have PPCS three months following injury ^5^. There are several PPCS phenotypes with the commonest symptoms being headaches, fatigue, sleep disturbances and cognitive difficulties ^6^. These symptoms have a detrimental impact on health-related quality of life and long-term outcomes ^7-9^.

Changes in the anatomy and function of whole-brain networks have shown to be associated with abnormalities in sleep regulation, fatigue and cognition ^10, 11^. Evidence suggests that the default mode network (DMN), which interlinks remote regions such as the posterior cingulate and medial prefrontal cortex, is critically impacted by TBI. Several studies have suggested that the DMN provides a sensitive indicator of the brain’s structural and functional integrity ^12-14^. For example, in stroke ^15^ and TBI ^11, 16, 17^, the anatomy and function of the DMN are compromised and alterations within this network have been linked to poor outcomes. DMN function has indeed been shown to reflect TBI-induced impairments in cognitive flexibility, sleep, and levels of fatigue ^11, 18^. In adults, TBI has also been linked to significant changes in the structure of the DMN ^17^. In line with these findings, structural changes in white matter pathways connecting DMN regions following mTBI have been linked to corresponding decreases in functional connectivity ^19^. DMN functions following TBI have also been attributed to deficits in sustained attention, wherein activity within the DMN is inconsistently regulated ^16^. Broadly, functional associations of DMN regions in other systems such as pain^20^, mood ^21^ and interoceptive awareness ^22^, could suggest a pervasive role for this network in behaviors commonly encountered in PPCS. Despite these current evidences, the converging impact that mTBI has on the structure and function of pediatric brains is unclear. Moreover, the association between brain changes linked to mTBI, PPCS, and clinical outcomes remains to be proven. On the basis of existing evidence, individual prediction of PPCS recovery may be achievable by examining changes in brain anatomy and function of the DMN following mTBI/concussion ^8^.

This report investigates a large prospective sample of children with established PPCS following mTBI. We started by examining whether acute post-injury structural and functional brain changes were linked to PPCS. By adopting an exploratory multivariate analysis, our previous study in the same sample identified key associations between *global* brain network measures of resting-state functional connectivity and PPCS. In line with existing findings ^17^, our previous results showed that a decrease in global DMN resting-state functional connectivity corresponded to increased sleep disturbances and cognitive difficulties in children with mTBI ^23^. Here, we expand on our prior findings which suggested that global patterns of functional connectivity (FC) were linked to sleep problems and fatigue in children following mTBI. In this study, we specifically tested the hypothesis that a whole-brain analysis of structure and function, associated to sleep disturbance and fatigue, converge to key regions and connections within the DMN. We employed local anatomical (voxel-based morphometry, VBM) and functional (regional homogeneity (ReHo) measures. These measures were complemented with the analysis of FC between regions showing significant brain-behavior associations. We also posit that these brain indices allow a significant and accurate prediction of clinical and cognitive recovery following mTBI.

## MATERIALS AND METHODS

### Subjects

We included 132 children with a confirmed diagnosis of mTBI: 99 children had PPCS at 4 to 6 weeks post-injury (henceforth referred as 1-month post-injury) and 33 children who had mTBI at 1-month post-injury but had recovered. We recruited a convenience sample provided by a sub-study of a clinical trial investigating the efficacy of melatonin in PPCS (PlayGame Trial: NCT01874847), conducted at the Alberta Children’s Hospital ^24^. The current study aimed to identify neuroimaging biomarkers of recovery in children following mTBI. There was a two-stage consent process. Participants with a medically diagnosed mTBI (defined using the American Academy of Neurology criteria ^25^) were recruited through the Emergency Department and gave consent to follow-up at 4 weeks post-injury. At this time, a clinical interview and examination was performed by a physician experienced in concussion assessment where proper consent was obtained. We applied the following exclusion criteria: (i) a previous concussion within 3-months; (ii) a more moderate to severe head injury (e.g. Glasgow Coma Scale (GCS) less than 13); iii) significant medical or psychiatric history; (iv) medications that likely affect participation in neuroimaging and/or sleep; and (v) inability to complete questionnaires and/or neuropsychological evaluation. The final cohort thus included 110 children (8.5 years to 17.9 years old) recovering from mTBI. None of the participants were taking any study medications at the time of imaging. For a comparative view with our neuroimaging measures (see *Analysis approach* section), we also collected data in an age-matched control group (*n* = 20, mean age = 14.44 years (standard deviation = 3.0 years), with 9 males and 11 females, see Table 1 in Iyer et al. (2019)^23^ for further demographic details). Ethical approval was granted by the University of Calgary (Canada) and the University of Queensland (Australia). Written consent from parents and child assent was obtained for all participants.

**Table 1.** Subject demographics and neuroimaging summary characteristics. A summary of sample size, age and gender distribution for the two clinical groups at 1-month post-injury (recovered and symptomatic PPCS); with median and standard deviation (SD) where appropriate. Gender and age distribution in each mTBI group, split by median age, is also provided. Regarding symptoms, the median symptom scale and PCSI (Likert-scale) for the whole cohort are shown. CNS VS: Computerized Neurocognitive Software Vital Signs, with the Neurocognition index as a standard score. Anatomical measures of total intracranial volume, grey matter (GM) and white matter (WM) reported across groups. Quality control measurements (FD) from rs-fMRI was also examined. The Chi Square test was used to assess putative group difference in Age whereas the Wilcoxon rank sum test was used to assess differences in symptoms between asymptomatic and symptomatic groups. “—” indicates non-significant values, * p_FDR_< 0.05, ** p_FDR_ < 0.001, controlling for false discovery rate (FDR).

| Sample characteristics |  | Statistics |  |  |  |
| --- | --- | --- | --- | --- | --- |
| *Demographic* |  | *mTBI*<br>(*n*=99) | *Recovered*<br>(*n*=31) | *Symptomatic*<br>(*n*=68) | *Significance* |
| Age |  | 14.5 (2.4) | 14.6 (2.24) | 15.12 (2.3) | — |
| Gender | *Male / female* | 48/51 | 21/10 | 27/41 | — |
|  | *<14.5 years* | 28/18 | 12/3 | 16/15 | — |
|  | *≥14.5 years* | 20/33 | 9/7 | 11/26 | — |
| *Symptoms* |  |  |  |  |  |
|  | *Sleep* | 1.80 (0.9) | 1.15 (0.3) | 2.21 (0.92) | ** |
|  | *Total PCSI* | 18.2 (0-75) | 4.90 (8.2) | 28.34 (17.1) | ** |
| ***Behavioral measures*** |  |  |  |  |  |
| CNS VS | *Neurocognition index (NCI)* | 98.20 (11.4) | 102.68 (7.8) | 97.13 (13.3) | * |
| ***Anatomical measures*** |  |  |  |  |  |
|  | Estimated total intracranial volume (TICV, cm<sup>3</sup>) | 1470.60 (132) | 1487.80 (129) | 1462.70 (135) | — |
|  | Grey Matter (GM, cm<sup>3</sup>) | 833.60 (83) | 851.30 (91) | 825.60 (79.5) | — |
|  | White Matter (WM, cm<sup>3</sup>) | 432 (54.6) | 434.80 (51) | 430.70 (57) | — |
| ***Functional measures*** |  |  |  |  |  |
|  | Frame-wise displacement, FD Power (mm) | 0.15 (0.04) | 0.14 (0.05) | 0.15 (0.04) | — |

#### Clinical and behavioral measures

We considered the following clinical and behavioral measures: (i) Post-Concussion Symptom Inventory (PCSI, parent proxy and child reports), which are standardized 26-item questionnaires used to summate key domains of post-concussive symptoms and behaviors ^26^. Specifically, total sleep scores were estimated using a sub-domain of the PCSI that assessed a subject’s level of drowsiness, less/more sleep than usual, trouble falling asleep, and increased fatigue; and (ii) a computerized cognitive assessment battery, CNS Vital Signs. Here the Neurocognition Index (NCI) was used as an overall performance index of cognitive and attentional processes ^27^. Subject demographics are summarized in **Table 1**.

Recovery was assessed at 4-6 weeks (time point, TP1) and 8-10 weeks post-injury (TP2) using overall change on the PCSI and a clinical interview conducted by a physician. Based on this interview and PCSI measurement, participants were then categorized into ‘*Symptomatic’* and ‘*Recovered’* groups. Children were considered as ‘*Symptomatic’* if there was a 10-point or greater increase in total PCSI score compared to pre-injury level. Participants were considered as ‘*Recovered*’ when their PCSI score returned to pre-injury level (see ***Supplementary Figure 1***, for study CONSORT diagram) ^28^. We here refer to asymptomatic status as to those children that *recovered* following mTBI. Following data quality control (see below), we included 69 children in the ‘*Symptomatic*’ group and 31 children in the ‘*Recovered*’ group. A machine learning classifier was trained to predict these dichotomous outcomes, while a complementary support vector machine (SVM) regression was used to assess the possibility of predicting outcome along a continuum (i.e. change in PCSI and NCI scores from 4 to 8-10 weeks post-injury). Processing and analyses of neuroimaging (provided below) was conducted on imaging data collected at TP1.

#### Neuroimaging

Images were acquired in the oblique axial plane using a 3.0 T GE scanner (Healthcare Discovery MR, 750w, 32-channel head coil). A structural scan (T1) was acquired using the following parameters: 0.8 mm slice thickness, flip angle = 10°, inversion time = 600 ms, FOV = 240 mm. Visual inspection of each T1 scan by a radiologist and/or neurologist excluded obvious signs of contusion or bleeding in grey and white matter. Resting-state functional magnetic resonance imaging (rs-fMRI) was recorded for five minutes and ten seconds via gradient echo planar imaging (scanning parameters included: EPI, TE = 30 ms, TR = 2000 ms, flip angle = 90, FOV = 230 mm, 64 × 64 matrix, slice thickness = 3.6 mm). The rs-fMRI acquisition time achieved a trade-off between the need for sufficient data and the necessity of minimizing movements and burden for unwell children ^29^. As per described in our recent report ^23^, head motion (Friston 24), linear trends, and signals from cerebrospinal fluid and white matter were removed using a general linear model framework. Data was bandpass filtered between 0.01 to 0.1 Hz. We excluded 10 subjects with less than 95% of data remaining after the removal of contaminated volumes (FD > 0.4 mm, including one preceding and two following volumes were censored). We also excluded 1 child due to incomplete clinical measures. Thus, we excluded 11 children from a cohort of 110, yielding a final sample size of 99 children (48 males and 51 females, no significant differences in gender). A summary on sample size characteristics and preliminary neuroimaging pre-processing details are provided in **Table 1**. As we used the same control cohort in our previous study ^23^, we did not exclude any healthy controls. **Table 1** provides an additional summary of neuroimaging characteristics in our final sample size.

#### Analysis approach

We started by testing for an association between structural (voxel-based morphometry, VBM) and functional (regional homogeneity, ReHo; functional connectivity, FC) brain indices with total sleep scores from the PCSI at 1-month post-injury. An overview of these initial analyses is shown in **Fig. 1**. Machine learning approaches (support vector machine, SVM) were subsequently implemented in an independent group to assess the ability of the selected brain indices to predict recovery outcomes that were unbiased by total sleep scores collected at TP1. Critically, for our SVM approaches we excluded total sleep scores from the PCSI inventory. Indices of total sleep score, however, were not significantly correlated with changes in cognitive (NCI) scores adopted for the regression SVM (*r* = 0.13, *p* = 0.18).

**Figure 1.**
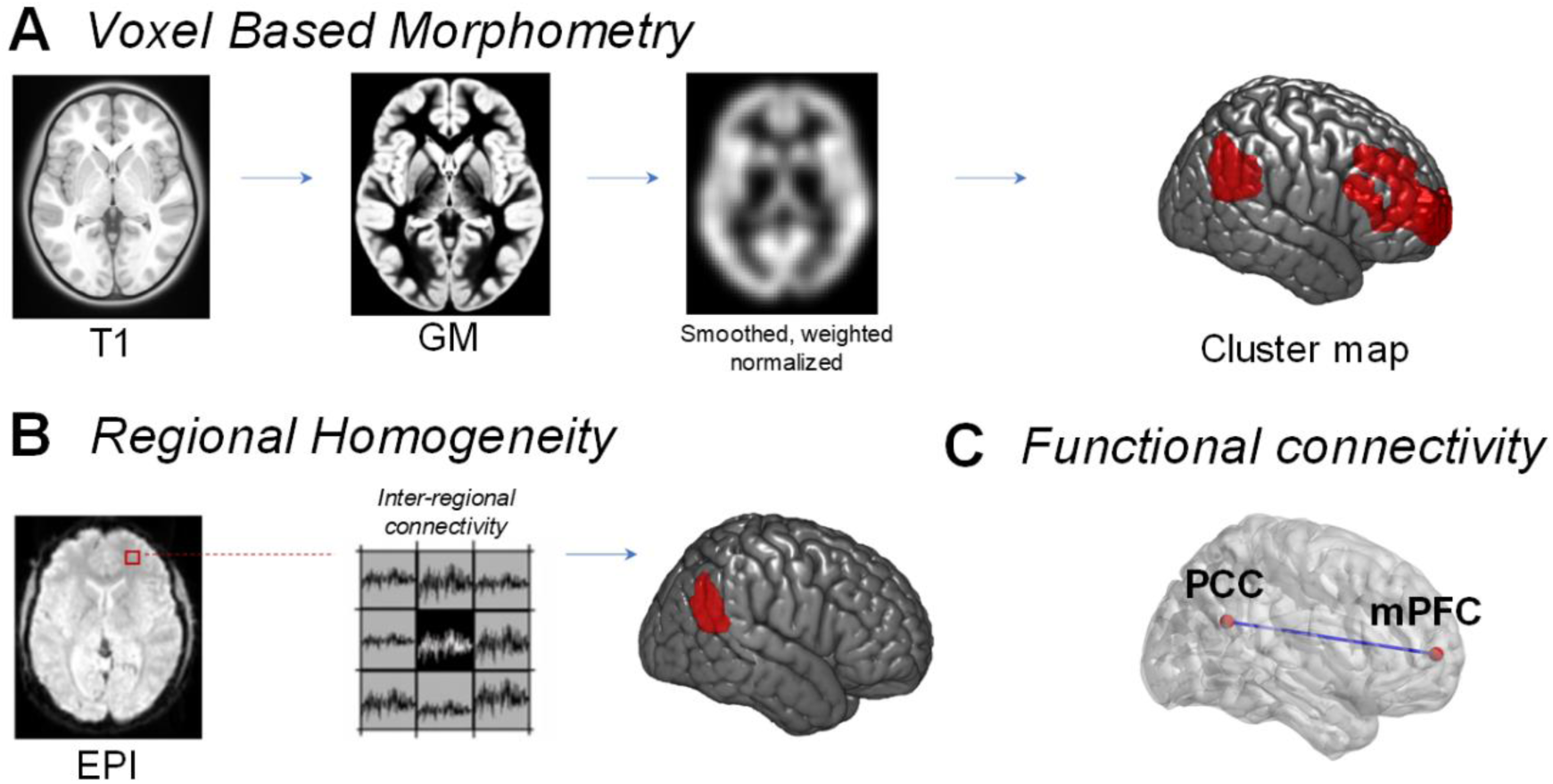
Processing pipeline for structural and functional data. **A** Voxel based morphometry (VBM) was performed using structural T1 scans from each child. Customized age-matched templates were used to derive GM, CSF and WM segmentations, before images were smoothed, weighted and normalized to MNI space. These normalized images were then used to assess the association between grey matter volume and a clinical construct assessing sleep disturbances. **B** Regional homogeneity (ReHo) assessment of functional imaging data (blood-oxygen-level-dependent, BOLD, data from echo planar imaging, EPI) was used to derive intra-regional connectivity within a set of neighboring voxels (n = 27). **C** Inter-regional functional connectivity was calculated using Pearson’s correlation between resting state BOLD signals in two brain regions of interest (posterior cingulate cortex (PCC) and medial prefrontal cortex (mPFC)).

We adopted an SVM approach to predict recovery outcomes, as this method offers several advantages over simpler methods including linear regression: (i) robustness to overfitting, particularly with high dimensional spaces, (ii) the use of Gaussian kernel scaling functions, which enables optimized thresholding of non-linear decision boundaries. This is critical for later assignment of decision labels (e.g. classifying *Recovered* versus *Symptomatic*), and (iii) appropriate margin of error in edge cases for labeling decisions (i.e. if a subject has opposing effects of one brain measure over other measures, SVM will maximize its decision margin to minimize error). Using a train-test approach, we trained our brain indices within an SVM classifier on 85% of subjects and tested the classifier’s accuracy on the remaining held-out individuals. This splitting of groups is in line with standard approaches in SVM classification ^30^, where data are “trained” on a larger group (usually between 80% - 90% of the sample) and “tested” on a smaller number of individuals. The selection of brain indices and training of the classifiers were therefore based on a sample of 85 individuals, with 14 individuals used as a naïve test sample to establish classification accuracy. In other words, the 14 individuals used to establish the trained classifier’s accuracy were not used to examine associations with our clinical variables, nor guide the selection of brain indices or classifier training. Classification was repeated 10 times (10-fold cross validation), with different sub-groups comprising a similar ratio of symptomatic and recovered individuals (*n =* 85 and *n =* 14; classifier results summarized in the **Supplementary Materials**). All preprocessing and analyses were performed within MATLAB (MathWorks, Natick, Massachusetts), using SPM12, DPARSFA toolboxes (V4.4) and custom scripts. Rendered 3D brain visualizations for our figures were generated using MRIcroGL.

##### VBM

A T1-weighted scan and VBM were used to define morphological markers ^31^, **Fig. 1A**. This was achieved by spatially normalizing all T1 images to the same stereotactic space and segmenting brain volumes into grey matter (GM), cerebrospinal fluid (CSF) and white matter (WM) images. A critical consideration for VBM in our study was the broad neurodevelopmental range (children aged between 8.5 and 18 years). We optimized VBM processing to account for variations in grey matter volume, brain size, and cortical thickness that are commonly associated with middle childhood and adolescence ^32^. Prior to segmentation of brain volumes, we first created customized tissue probability maps (TPMs), to appropriately reflect age and gender differences. Specifically, our TPM maps were created via the use of pediatric neuroimaging templates available from the NIH MRI study of Normal Brain Development ^33-35^. The age range (asymmetric templates, 4.5 to 18.5 years) and features present within these template images were chosen to appropriately reflect natural variations of cortical thickness and grey matter over the course of childhood ^36^. To generate these TPMs we used the Template-O-Matic (TOM8) toolbox ^37^, so that age groups could be segmented with appropriate TPMs-i.e. a TPM could be allocated for each year between 8 and 18 years.

Using age-appropriate TPMs, we next segmented brain volumes into grey matter, CSF and WM via the SPM12 toolbox using default segmentation parameters and affine regularization with an average sized template to maximize spatial accuracy within age-groups. A data quality check was performed on all brain volumes. Next, diffeomorphic anatomical registration ^38^ was performed to iteratively model brain shape differences arising from GM and WM images and optimize accuracy of inter-subject alignment whilst correcting for volume changes. All images were then spatially normalized to derive smoothed (modulated), weighted, Jacobian scaled grey matter images localized to the MNI space. We employed an isotropic Gaussian kernel at 12 mm full-width at half maximum (FWHM) for smoothing, whilst specifying a voxel size of 3 mm for spatially normalized images. As per our segmentation approach, we performed image normalization within specific age-groups (e.g. 8 to 9 years, 9 to 10 years etc.) to limit false positive errors that may be generated in larger group sizes ^39^. Smoothed images were checked across age groups for consistency (N, median = 11 subjects, across 10 age-groups). Prior to statistical analysis, all grey matter volumes were then processed with an absolute threshold mask of 0.2, to censor the possible influence of edge-effects present in other tissues (e.g. white matter).

A general linear model (GLM) was used to examine the relationship between grey matter volume and total sleep score. Results were corrected for family-wise error at cluster-level (*p*_FWE_ < 0.05, search threshold *p*_uncorr_ < 0.001, cluster-level threshold (*k*_E_) > 500). Statistical tests were also re-performed on smoothed images processed at a smaller kernel size (FWHM, 9 mm) with the same family-wise error correction parameters to validate our findings ^39^.

##### ReHo (*intra*-regional connectivity)

Processing of rs-fMRI data is specified in our recent report ^23^. Importantly, for ReHo analysis described herein unsmoothed data were used. Briefly, ReHo analysis provides a validated metric of local intra-regional functional connectivity ^40^ calculated by assessing the temporal synchronization (*or* also known as Kendall coefficient of concordance, KCC) of the blood-oxygen-level-dependent (BOLD) signal between a small (*n* = 27)^41^ set of spatially adjacent voxels (**Fig. 1B**). In this study, whole-brain ReHo estimates provide a multimodal measure that links local VBM estimates with between region functional connectivity. Whole-brain ReHo estimates were used to assess brain-symptoms relationships (*p*_FWE_ < 0.05 at cluster-level, search threshold *p*_uncorr_ < 0.005).

##### FC (*inter*-regional connectivity)

In our prior study ^23^, we adopted a whole-brain parcellation ^42^ to examine changes in functional connectivity present within canonical resting-state networks in children with mTBI. In this study, guided by our VBM and ReHo results, we utilize a seed-based approach for measuring inter-regional FC (**Fig. 1C**). We placed two spherical seeds within the brain right-hemisphere: (i) the posterior cingulate cortex (PCC, MNI coordinates: *x* = 15 mm, *y =* −57 mm, *z* = 24 mm; 8 mm radius sphere) and (ii) the medial prefrontal cortex (mPFC, MNI coordinates: *x* = 11 mm, *y* = 58 mm, *z* = 6 mm, 8 mm radius sphere). All subjects’ Pearson correlation coefficients (*r*) were calculated using BOLD timeseries from the PCC and mPFC. The resulting *r*-values were subsequently Fisher-Z transformed. Individual FC values were subsequently used to perform correlations with our clinical variables.

##### Machine learning

We next sought to investigate the utility of the structural and functional brain indices to predict follow-up measures of cognition and recovery. SVM regression ^43^ was utilized to determine the ability of the selected brain indices (VBM, ReHo, and FC) to predict change in PCSI and NCI scores.

A SVM classifier ^43^ was also performed on 85 children to determine if brain indices of interest could be used to discriminate recovery from dichotomous outcomes of PPCS (recovered *vs*. symptomatic). SVM classification was used to predict dichotomous clinical outcomes via a hold-out cross-validation (*n* = 14 used as test sample). The SVM accuracy was further tested using 10-fold cross-validation in 9 additional data groupings (i.e. each additional group comprised 85 individuals as training samples and 14 individuals as naïve test samples). The average performance of both SVM regression and SVM classifier results are presented in the Results section.

## RESULTS

### DMN structure and function in children with PPCS is related with sleep disturbances

We found a significant negative association between PCC and mPFC grey matter volumes and sleep scores: Grey matter volume in these brain regions was lower in children with higher sleep disturbances (*n =* 85, *r =* −0.30 ± 0.008 standard error (S.E.) across 10-folds, *p*_FWE_ = 0.002 for PCC, *r* = −0.38 ± 0.013 S.E., *p*_FWE_ = 0.0001 for mPFC; **Fig. 2A**). Negative associations between grey matter volumes and total sleep scores were also confirmed for spatially smoothed at a kernel size of 9 mm (across 10-folds, *r* = −0.34 ± 0.012 S.E. for PCC *p*_FWE_ = 0.0013, *r* = −0.14 ± 0.02 S.E., *p*_FWE_ = 0.0003 for mPFC).

**Figure 2.**
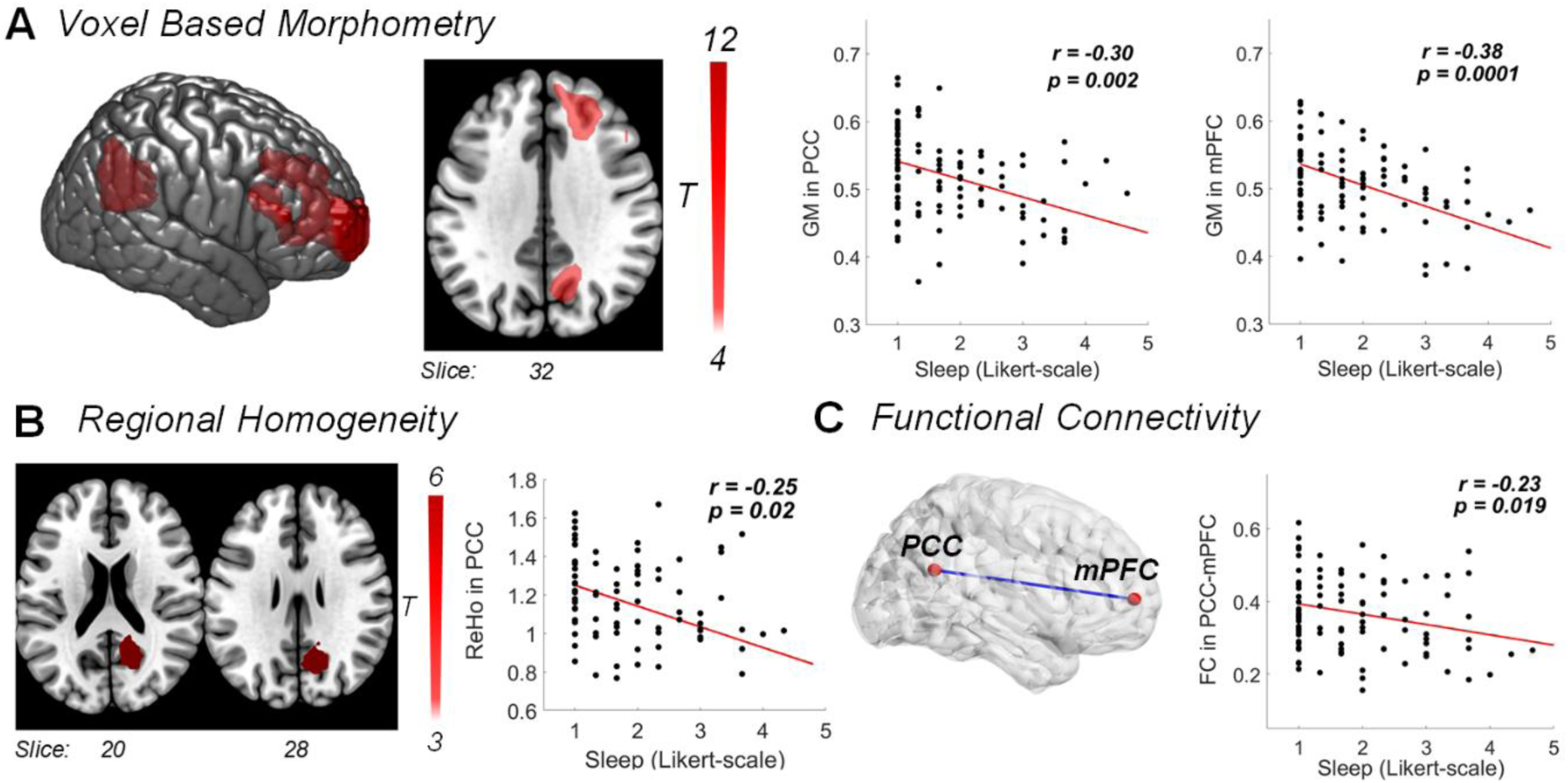
Association between brain - structure and function - and symptoms. **A** Grey matter volume changes, detected using voxel-based morphometry (VBM), negatively correlated with sleep scores such that decreased grey matter volume in the right PCC and the right mPFC was linked to increased sleep disturbances and fatigue (cluster-level *p*_FWE_ < 0.002 and *p*_FWE_ < 0.0001, respectively; high threshold of *p*_uncorr_ < 0.001). **B** Reduced within-region functional connectivity (ReHo) in the right PCC also negatively correlated with sleep problems (*p*_FWE_ = 0.02 at cluster-level, high threshold of *p*_uncorr_ < 0.005). **C** Functional connectivity (FC) between the right PCC and mPFC indicates a negative correlation: Children with lower across-region FC showed increased sleep problems. Results in this figure are from the first data grouping (fold) 1. All correlations were adjusted for age, gender and brain volumes.

ReHo in the PCC was negatively associated with sleep problems (*r =* −0.25 ± 0.036 S.E., *p*_FWE_ = 0.02; **Fig. 2B**). In line with the aforementioned grey matter volume associations, decreased ReHo in the PCC was related to increased sleep problems.

In agreement with the associations between local brain indices and sleep-related scores, FC *between* the PCC and mPFC also negatively correlated with sleep scores (*r =* −0.23 ± 0.0081, *p* = 0.019; **Fig. 2C**). Across our measures, we only observed an effect of gender for the correlation between measures of FC between the PCC and mPFC and sleep scores (female *r* = −0.28 ± 0.02 S.E., *p* = 0.006, male children *r* = −0.08 ± 0.03 S.E., *p* = 0.61). Following this observation, we investigated possible differences in sleep symptom severity between genders. Results showed higher sleep disturbances in females compared to males (unpaired t-test, t_97_ =3.20, *p*=0.002). However, values of FC were similar between genders (unpaired t-test, t_97_ =1.55, *p*=0.17). See ***Supplementary Table 1*** for a summary of associations and the relative contribution of age, gender and brain volumes.

Additionally, the comparison between brain indices of mTBI children with our age-matched controls revealed group differences in grey matter and ReHo measures, but not in FC (one-way ANOVA we report significant differences for GM in PCC: *p*_FDR_ = 1.1 × 10^−12^; GM in mPFC: *p*_FDR_ =1.2 × 10^−9^; ReHo in PCC: *p*_FDR_ = 5.3 × 10^−7^; FC: *p*_FDR_ =0.25, see ***Supplementary Figure 2*)**.

### Brain signatures of sleep carry prognostic information

SVM regression included the four brain indices collected at 1-month post-injury: grey matter volume (eigenvariates) extracted from the PCC and mPFC, ReHo estimates from the PCC, and PCC-mPFC FC values. Age at 1-month post-injury was also included. Changes in PCSI scores (8-10 weeks follow-up scores *minus* 4 weeks post-injury scores; total sleep scores excluded) predicted by these five indices were highly correlated with real PCSI scores (10-folds average: *r* = 0.52, *p* = 0.009, 12.28% standard error; **Fig. 3A**). Moreover, SVM regression using the same brain indices was also able to predict changes in cognitive (NCI) scores (10-folds average: *r* = 0.43, *p* = 0.0002, 4.40% standard error).

**Figure 3.**
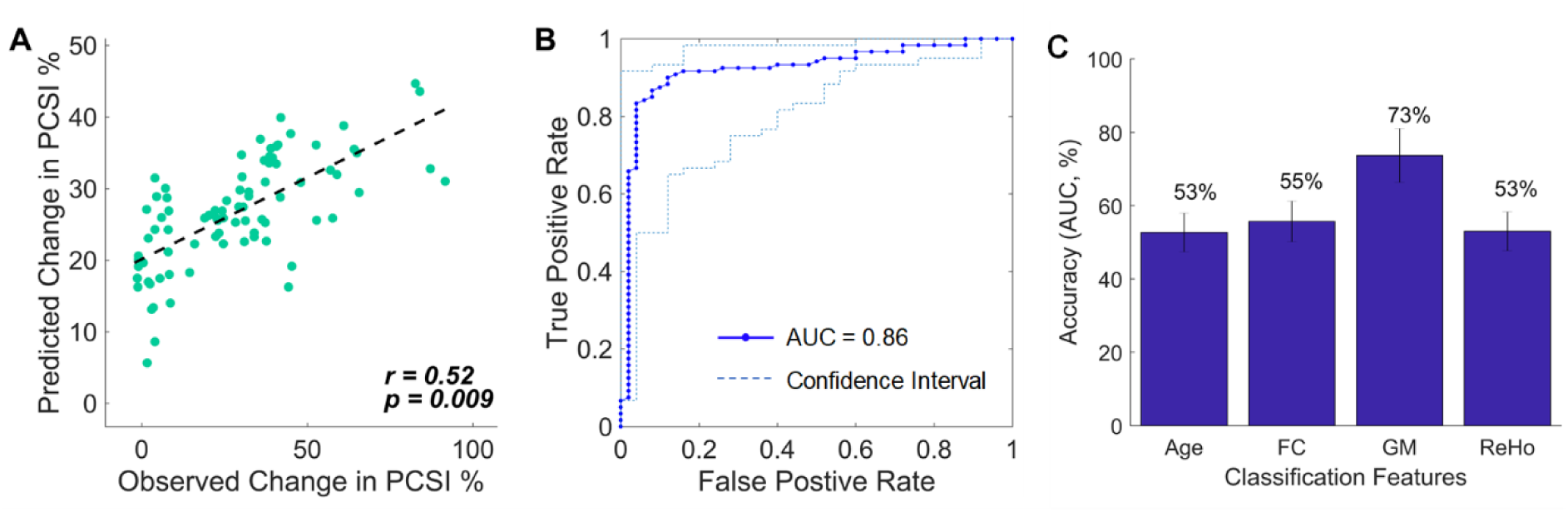
Classification and outcome assessment. **A** SVM regression was performed using four brain features linked to behavioral sleep scores at 1-month post-injury: PCC and mPFC grey matter volumes, PCC values of local functional connectivity (ReHo), and resting-state functional connectivity between PCC and mPFC. These brain features, plus the age of the participant, were able to significantly predict changes in post-concussion symptoms between 1-month post-injury and 8-10 weeks follow-up along a continuum. PCSI = Post-Concussion Symptom Inventory. **B** SVM classification results using the four brain features of interest and age of the child to classify recovery *versus* symptomatic PPCS at follow-up (8-10 weeks mark). The receiver operating characteristic curve (ROC) shows an area under the curve (AUC) with high specificity and sensitivity. The confidence interval (dashed lines) indicates the most and least accurate classification across the 10-folds. **C** Accuracy of age and each brain feature (GM, ReHo and FC) - tested independently - in predicting recovery outcome. Error bars indicate averages over 10-folds (± standard deviation).

The SVM classifier, adopting the abovementioned four brain features plus age, was able to accurately discriminate children that recovered from concussion to those that remained clinically symptomatic (10-folds average: 86% accuracy, 69% specificity and 94% sensitivity; **Fig. 3B**). The accuracy range for the classifier to discriminate *symptomatic* from *recovered* children was between 78% to 92% (10-folds; **Fig. 3B**). The prediction accuracy on our hold-out test samples (*n* = 14) also maintained high accuracy (10-folds average: 79% accuracy, 75% specificity, and 82% sensitivity). The range of accuracy in the test samples was between 65% to 95% (10-folds). See **Supplementary Table 2 and 3** for a summary of classification results.

## DISCUSSION

The current study supports the notion that structural and functional brain changes in the DMN underpin core manifestations of pediatric PPCS following mTBI ^44^. Our findings also extend upon previous work by showing that indices of symptom-related changes in grey matter and functional connectivity within and between two key regions of the DMN — PCC and mPFC — carry significant prognostic information.

Prior studies in both children and adults have suggested that the DMN plays a crucial role in sleep processes ^45^, which in turn helps maintain cognitive performance ^46, 47^. Accordingly, it has been shown that TBI causes structural and functional changes within the DMN that have been linked to cognitive deficits, including slower rates of information processing and decision-making ^17, 18, 48, 49^. Though the neural underpinning of poor sleep and excessive fatigue following mTBI has not been elucidated by prior work, recent studies in adolescents have pointed towards weaker DMN functional connectivity ^50^. Our study directly supports and extends these emerging evidences by showing that converging decreases in grey matter volume, as well as intra and inter-regional functional connectivity, in the PCC and mPFC in children with mTBI-induced PPCS are linked to increases in sleep disturbances and fatigue. Negative associations of grey matter and intra-regional connectivity with sleep-related problems occurred within the context of higher brain index values compared to matched healthy controls (***Supplementary Figure 2***). These findings provide support to the notion of a neurotrauma-related compensatory response within DMN regions ^18^. Although grey matter and intra-regional connectivity values did not indicate gender-specific associations, the negative association between inter-regional FC and sleep disturbances was driven by females. This association, along with the fact that females reported higher levels of sleep-related symptoms compared to males, is in line with the suggestion that concussed females are more sensitive to sleep difficulties and have protracted recovery patterns ^8, 51^.

Cognitive functions, including reasoning and working memory, have been associated with higher DMN functional connectivity at rest ^52, 53^. Accordingly, poorer cognitive functions following TBI have been linked to reduced DMN structural and resting-state connectivity ^7, 17, 54, 55^. In our previous exploratory analysis conducted on the same cohort, we were unable to detect a significant change in *global* DMN resting-state functional connectivity in children following mTBI compared to matched controls ^23^. We did, however, find an association between variations in DMN global connectivity and cognitive symptoms. The current re-analysis of the data offers an important insight into our previous results, indicating that key symptoms of PPCS following mTBI are linked to circumscribed changes to the structure and function of two key DMN nodes. Our linear classifiers build upon this assertion, showing that a combination of PCC and mPFC structural and functional features can significantly and accurately predict cognitive outcomes. DMN functional connectivity in children has shown to normalize following improvements in cognitive functions ^56^ and sleep quality ^50^. Our findings are consistent with these previous reports and highlight that a normalization of functional interactions between, as well as within, the PCC and the mPFC are most likely responsible for the observed behavioral recovery.

Recovery from PPCS in children is complex to predict. Monitoring of symptoms and clinical assessments using scales like the PCSI have shown some sensitivity in predicting behavioral outcomes ^8^. However, the predictive value of psychometric and behavioral measurements in tracking recovery trajectory alone is insufficient, limiting the ability to develop and implement outcome-oriented personalized therapeutic interventions. Our findings motivate the use of neuroimaging indices to supplement existing clinical evaluations of symptoms and behaviors during recovery from PPCS. Specifically, the anatomical and functional assessment PCC and mPFC provide useful insights on the likelihood of recovery of a developing injured brain. In fact, our findings suggest that convergent anatomical and functional changes within and between these two DMN regions offer valuable prognostic information on the temporal evolution of core PPCS. These results echo prior neuroimaging studies showing that improvement in adult recovery following TBI is linked with higher activity within the DMN^17,19^.

Predicting the recovery trajectory of a child with PPCS is encumbered by the high inter-individual variability in symptoms, age, gender, and pre-morbid characteristics ^8^. Our findings showed that the inter-individual variability of PPCS recovery in children is linked to variations in brain indices that are indexed by relatively simple clinical scores of sleep disturbance and fatigue. In this context we note that, while appealing, a dichotomous classification of outcomes (recovered *versus* symptomatic) may misrepresent the actual progression of PPCS following mTBI. In effect, our machine learning results highlight that changes in PPCS including cognitive functions can be predicted by an SVM regression approach. That is, the collection and assessment of anatomical and functional brain indices alongside standard neuropsychological assessments at different timepoints post-injury offer potential for prognosis along a natural continuum of recovery. More broadly, our results are in agreement with the suggestion that recovery from mTBI maps onto a dynamic and “multifaceted construct” ^8^.

In sum, our study complements prior work in supporting the key role of sleep and fatigue to the clinical picture of PPCS ^23, 44^. The significant association between variations in such key symptoms and structure-function in PCC and mPFC directly support the notion that the DMN underpins core manifestations of mTBI-induced PPCS. The prognostic value of these brain indices further encourages this hypothesis and provide a strong rationale for translational efforts aiming to facilitate the use of neuroimaging in clinical settings. Future validation studies on independent datasets are required to test and improve the predictive utility of the brain indices identified in this study towards overall recovery outcomes. The provision of additional information gathered from neuroimaging may indeed enable the targeted allocation of limited therapeutic resources to children with the greater likelihood of poor outcome.

## ACKNOWLEDGEMENTS

K.K.I acknowledges funding support from the Brain Foundation. K.M.B. acknowledges funding support from the Motor Accident Insurance Commission (MAIC, Queensland, Australia) and the Canadian Institutes of Health Research (grant number: 293375). L.C. is supported by the Australian National Health Medical Research Council (L.C. 1099082 and 1138711). We thank Dr Nathan Stevenson for his advice on implementation of support vector machine classification with our dataset.

## AUTHOR CONTRIBUTIONS

K.K.I. performed all aspects of pre-processing, analysis, interpretation of results, writing of the manuscript and evaluation of findings. A.Z. assisted with analysis, statistical inferences, writing of the manuscript and evaluation of findings. K.M.B. conceived, designed, acquired data for the study and provided clinical interpretation of the findings. L.C. performed all aspects of pre-processing, analysis, interpretation of results, writing of the manuscript and evaluation of findings.

## COMPETING INTERESTS

No competing interests or conflicts of interest to declare.

## Supplementary Materials

This supplementary material has been provided by the authors to give readers additional information about their work.

**Supplementary Figure 1.**
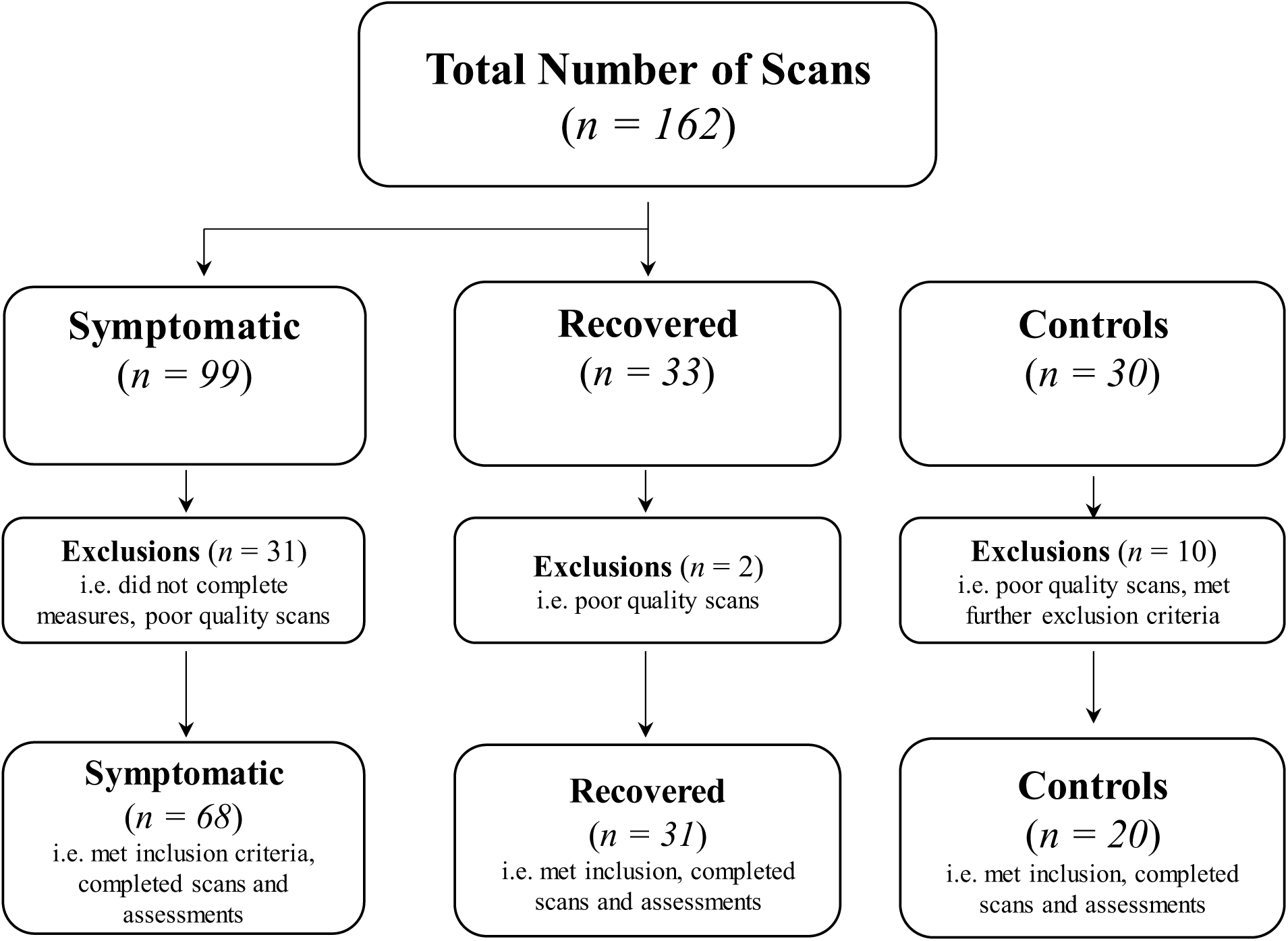
Study CONSORT diagram. For this study, there were a total of 99 mTBI children and 20 healthy controls who met study inclusion criteria, and passed quality control measures following pre-processing of neuroimaging scans.

**Supplementary Table 1.** General linear modelling results. The table summarizes correlations for grey matter (GM), regional homogeneity (ReHo) and functional connectivity (FC), following adjustment for age, gender and total intracranial volume (TICV). A summary of each variable’s t-statistic, statistical significance with standard error observed across 10-folds is provided.

| Measures | T-statistic | Significance (p-value) | Standard error |
| --- | --- | --- | --- |
| *Anatomical (GM)* |  |  |  |
| *PCC* |  |  |  |
| Age | 2.37 | 0.02 | 0.11 |
| Gender | 1.88 | 0.06 | 0.58 |
| GM | -2.86 | 0.01 | 4.82 |
| TICV | 0.52 | 0.61 | 2.33 |
| *mPFC* |  |  |  |
| Age | 2.57 | 0.01 | 0.11 |
| Gender | 1.68 | 0.10 | 0.56 |
| GM | -4.02 | 0.00012 | 4.79 |
| TICV | 0.62 | 0.54 | 2.17 |
| *Intra-regional FC (ReHo)* |  |  |  |
| *PCC* |  |  |  |
| Age | 2.55 | 0.01 | 0.11 |
| Gender | 1.44 | 0.15 | 0.58 |
| ReHo | -3.38 | 0.0011 | 1.15 |
| TICV | -0.58 | 0.56 | 2.10 |
| *Inter-regional FC* |  |  |  |
| *PCC-mPFC* |  |  |  |
| Age | 2.06 | 0.041 | 0.04 |
| Gender | 2.47 | 0.02 | 0.20 |
| FC | -1.99 | 0.048 | 0.82 |
| TICV | 0.68 | 0.50 | 0.75 |

**Supplementary Figure 2.**
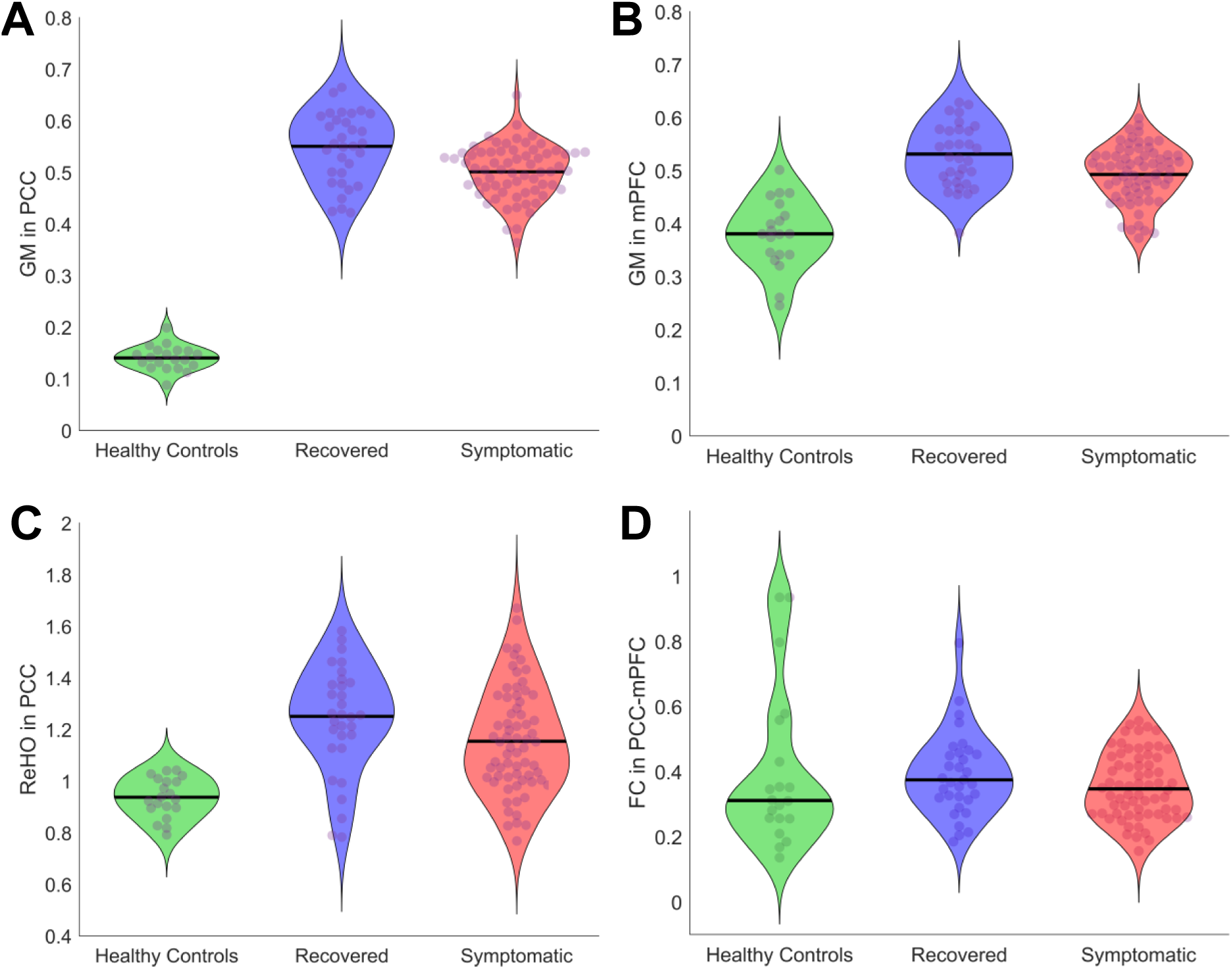
Brain indices in mTBI and age-matched healthy controls. **A-B** Grey matter volume (eigenvariate) measures in PCC and mPFC, respectively. **C** Regional homogeneity (ReHo) values measured in the posterior cingulate cortex (PCC). **D** Functional connectivity values between the PCC and the mPFC. Black lines on violin plots indicate median values. Individual subjects are represented by dots via a beeswarm plot. Demographic details on age-matched controls are provided in our previous report^23^, see Table 1 for the sample size characteristics of healthy controls.

**Supplementary Table 2.**
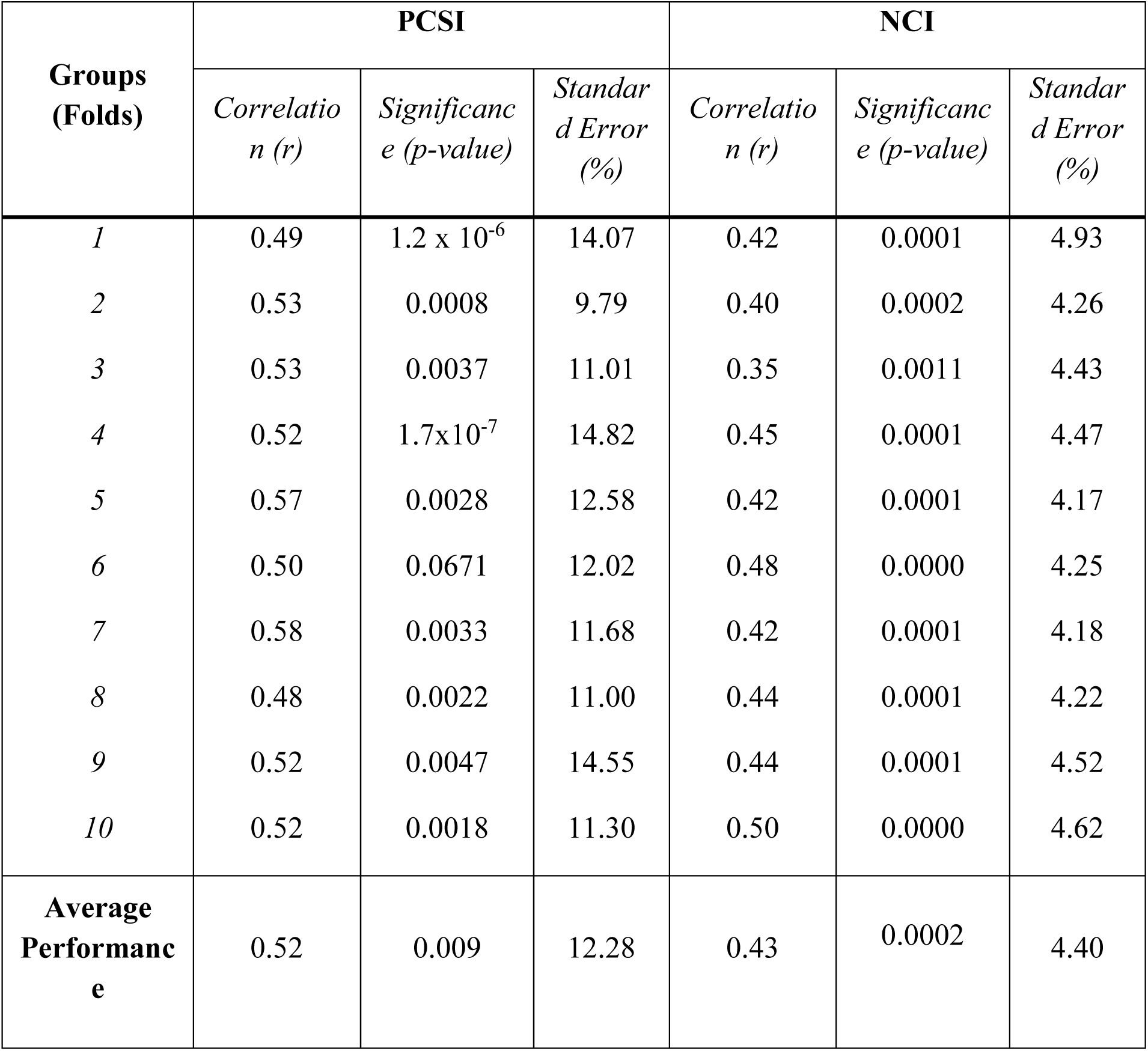
SVM regression results across 10 different groups (folds). The table summarizes the relation between predicted (using brain indices) and observed changes in clinical scores. Specifically, the table summarize Pearson’s correlation coefficient (*r*), significance (*p-value)* and standard error (%) across 10 folds (training group *n* = 85, test group *n* = 14).

| Groups (Folds) | PCSI |  |  | NCI |  |  |
| --- | --- | --- | --- | --- | --- | --- |
|  | Correlation (r) | Significance (p-value) | Standard Error (%) | Correlation (r) | Significance (p-value) | Standard Error (%) |
| 1 | 0.49 | $1.2 \times 10^{-6}$ | 14.07 | 0.42 | 0.0001 | 4.93 |
| 2 | 0.53 | 0.0008 | 9.79 | 0.40 | 0.0002 | 4.26 |
| 3 | 0.53 | 0.0037 | 11.01 | 0.35 | 0.0011 | 4.43 |
| 4 | 0.52 | $1.7 \times 10^{-7}$ | 14.82 | 0.45 | 0.0001 | 4.47 |
| 5 | 0.57 | 0.0028 | 12.58 | 0.42 | 0.0001 | 4.17 |
| 6 | 0.50 | 0.0671 | 12.02 | 0.48 | 0.0000 | 4.25 |
| 7 | 0.58 | 0.0033 | 11.68 | 0.42 | 0.0001 | 4.18 |
| 8 | 0.48 | 0.0022 | 11.00 | 0.44 | 0.0001 | 4.22 |
| 9 | 0.52 | 0.0047 | 14.55 | 0.44 | 0.0001 | 4.52 |
| 10 | 0.52 | 0.0018 | 11.30 | 0.50 | 0.0000 | 4.62 |
| Average Performance | 0.52 | 0.009 | 12.28 | 0.43 | 0.0002 | 4.40 |

**Supplementary Table 3.** Outcome classifier (SVM) results across 10 different groups (folds). The table summarizes accuracy, sensitivity, and specificity values across 10 different folds (training group *n* = 85, test group *n* = 14). The average performance across the folds indicates a high accuracy in predicting recovery outcomes.

| Groups (Folds) | Accuracy |  | Sensitivity |  | Specificity |  |
| --- | --- | --- | --- | --- | --- | --- |
|  | Training | Test | Training | Test | Training | Test |
| 1 | 0.82 | 0.95 | 0.88 | 1 | 0.67 | 0.88 |
| 2 | 0.88 | 0.80 | 0.96 | 0.83333 | 0.70 | 0.75 |
| 3 | 0.88 | 0.95 | 0.96 | 1 | 0.70 | 0.88 |
| 4 | 0.88 | 0.80 | 0.96 | 0.83333 | 0.70 | 0.75 |
| 5 | 0.92 | 0.65 | 1 | 0.66667 | 0.72 | 0.63 |
| 6 | 0.88 | 0.65 | 0.96 | 0.66667 | 0.70 | 0.63 |
| 7 | 0.85 | 0.70 | 0.92 | 0.66667 | 0.68 | 0.75 |
| 8 | 0.85 | 0.80 | 0.92 | 0.83333 | 0.68 | 0.75 |
| 9 | 0.88 | 0.80 | 0.96 | 0.83333 | 0.70 | 0.75 |
| 10 | 0.78 | 0.80 | 0.84 | 0.83333 | 0.65 | 0.75 |
| **Average**<br>**Performance** | 0.86 | 0.79 | 0.82 | 0.82 | 0.70 | 0.75 |

## Notes

#### Summary of Updates

Supplemental files updated

